# *Geobacter sulfurreducens* extracellular multiheme cytochrome PgcA facilitates respiration to Fe(III) oxides but not electrodes

**DOI:** 10.1101/172775

**Authors:** Lori Zacharoff, Dana Morrone, Daniel R. Bond

**Author notes:** Current address: University of Southern California, 920 Bloom Walk, Los Angeles, CA 90034. **Send all correspondence to:** Daniel R. Bond 140 Gortner Laboratory 1479 Gortner Ave St. Paul, MN 55108.

## Abstract

Extracellular cytochromes are hypothesized to facilitate the final steps of electron transfer between the outer membrane of the metal-reducing bacterium *Geobacter sulfurreducens* and solid-phase electron acceptors such as metal oxides and electrode surfaces during the course of respiration. The triheme *c*-type cytochrome PgcA exists in the extracellular space of *G. sulfurreducens*, and is one of many multiheme *c-*type cytochromes known to be loosely bound to the bacterial outer surface. Deletion of *pgcA* using a markerless method resulted in mutants unable to transfer electrons to Fe(III) and Mn(IV) oxides; yet the same mutants maintained the ability to respire electrode surfaces and soluble Fe(III) citrate. When expressed and purified from *Shewanella oneidensis*, PgcA demonstrated a primarily alpha helical structure, three bound hemes, and was processed into a shorter 41 kDa form lacking the lipodomain. Purified PgcA bound Fe(III) oxides, but not magnetite, and when PgcA was added to cell suspensions of *G. sulfurreducens,* PgcA accelerated Fe(III) reduction similar to addition of FMN. Addition of soluble PgcA to ∆*pgcA* mutants also restored Fe(III) reduction. This report highlights a distinction between proteins involved in extracellular electron transfer to metal oxides and poised electrodes, and suggests a specific role for PgcA in facilitating electron transfer at mineral surfaces.

## Introduction

Dissimilatory metal reducing bacteria such as *Geobacter sulfurreducens* have to transfer respiratory electrons to extracellular acceptors via direct contact to minerals such as iron and manganese oxides. These minerals exist as a heterogeneous mixture of insoluble particles in nature, with a range of redox potentials and surface charges that change during reduction (Byrne et al., 2011; Coker et al., 2012; Cutting et al., 2009; Majzlan, 2012; Nealson and Saffarini, 1994). Respiration of such diverse acceptors in soils and sediments is likely to require continuous modification of the extracellular space to facilitate interfacial contact. Evidence is accumulating that *Geobacter* strains alter secretion of polysaccharides (Rollefson et al., 2011), conductive pili (Klimes et al., 2010; Reguera et al., 2005), and multiheme *c*-type cytochromes (Ding et al., 2006, 2008; Mehta et al., 2005; Nevin et al., 2009), depending on environmental conditions.

A few *G. sulfurreducens* proteins are known to be secreted beyond the outer membrane where they could act as loosely bound or mobile mediators to facilitate the final steps of electron transfer analogous to how secreted redox-active molecules accelerate reduction by *Shewanella oneidensis* (Lies et al., 2005; Marsili et al., 2008; Von Canstein et al., 2008), *Geothrix fermentans* (Mehta-Kolte and Bond, 2012; Nevin and Lovley, 2002) and *Geobacter uraniireducens* (Tan et al., 2016). For example, the tetraheme cytochrome OmcE can be physically sheared from intact Mn(IV) oxide grown cells (Mehta et al., 2005), the hexaheme cytochrome OmcS complexes with pili during growth with Fe(III) oxides (Leang et al., 2010), and the octaheme cytochrome OmcZ is associated with the extracellular matrix of cells grown on electrodes (Inoue et al., 2011). A more elusive secreted cytochrome was described in 1999 (Lloyd et al., 1999), where a 41 kDa extracellular hemeprotein enriched from *G. sulfurreducens* supernatants was found to rapidly adsorb to Fe(III) oxides. However, this protein has not been linked to a genetic locus.

One candidate for this uncharacterized 41 kDa extracellular cytochrome is the *c-*type triheme lipocytochrome PgcA (GSU1761). Expression of *pgcA* is driven by a GEMM (genes for the environment, membranes and motility) riboswitch responsive to the dinucleotide cyclic AMP-GMP (Kellenberger et al., 2015; Nelson et al., 2015). In proteomic surveys, PgcA is more abundant when insoluble Fe(III) oxides are the terminal electron acceptor, compared to soluble Fe(III) citrate (Ding et al., 2008). Expression of *pgcA* also increases during growth with Fe(III) oxide compared to Fe(III) citrate (Aklujkar et al., 2013). Selection for rapid growth with Fe(III) oxides enriches for riboswitch mutations that enhance *pgcA* expression, and selection of a *G. sulfurreducens* KN400 pili mutant for improved Fe(III) oxide reduction increased *pgcA* expression, and led to production of a ∼40 kDa extracellular cytochrome identified as PgcA. As the *pgcA* gene predicts a 57 kDa product, this result suggests processing of secreted PcgA by cells (Smith et al., 2014; Tremblay et al., 2011; Yi et al., 2009).

With a predicted localization as a lipoprotein on the cell surface, detection in a processed unbound form, and link to metal oxide reducing conditions, PgcA could play an unrecognized role in the final stages of extracellular electron transfer by *G. sulfurreducens* (Smith et al., 2014; Tremblay et al., 2011; Yi et al., 2009). Here, we investigated PgcA by creating and complementing markerless *pgcA* (∆*pgcA*) deletion strains, and purifying PgcA from a heterologous host. Mutants lacking *pgcA* were severely deficient in Fe(III) oxide respiration, but remained unimpaired in growth with other extracellular acceptors such as electrodes and soluble Fe(III) citrate. We found heterologous PgcA to exist in two forms: 57 kDa as well as a shorter 41 kDa domain lacking the predicted lipid attachment site. Purified PgcA bound Fe(III) oxides but not magnetite, and when added to resting cell suspensions of both wild type and ∆*pgcA* cultures, soluble PgcA accelerated Fe(III) reduction similar to added flavin mononucleotide. This implicates PgcA-family cytochromes as a class of proteins specific to metal oxide reduction.

## Materials and Methods

### Cell culture and growth assays

Laboratory stocks of *G. sulfurreducens* PCA (lab strain resequencing described in (Chan et al., 2015)), and mutants were resuscitated from laboratory stocks by streaking onto 1.5% agar containing minimal salts medium (NB) (Chan et al., 2015) with 20 mM acetate and 40 mM fumarate (NBFA), and picking colonies into liquid medium for each experiment. All *G. sulfurreducens* cultures and media were prepared anaerobically under 80% N_2_, 20% CO_2_ atmosphere.

Electrochemical bioreactor experiments contained NB medium with 20 mM acetate and additional NaCl salts added in place of fumarate. Cultures of *Geobacter* strains grown with excess acetate, were used as an inoculum as they approached an OD (600 nm) of 0.5. Polished graphite electrodes (1500 grit), with a surface area of 3 cm^2^, were used as working electrodes. A small piece of platinum wire was used as a counter electrode and a calomel electrode connected via a Vycor frit salt bridge was used as a reference electrode. Bioreactors were maintained at a constant 30 °C. Growth with freshly precipitated insoluble Fe(III) oxide (55 mM), Fe(III) citrate (55 mM) and MnOOH (30 mM) was performed in the same medium without the additional salt and 20 mM acetate as the electron donor.

Fe(III) reduction was measured by monitoring accumulation of Fe(II) by means of a FerroZine assay. As previously described, (Levar et al., 2014; Rollefson et al., 2009) 100 μL samples were extracted from 10 mL Balch culture tubes and diluted in 1 N hydrochloric acid until conclusion of the experiment when the FerroZine assay was performed in 96 well plate format. Mn(IV) reduction was monitored as described in (Levar et al. 2017).

### Biofilm formation

Cell attachment to surfaces was characterized using a crystal violet, 96 well plate assay as described previously (Rollefson et al., 2009, 2011). Growth medium contained 30 mM acetate, 40 mM fumarate. Incubation occurred for 72 hours. Optical density at 600 nm was measured, wells were emptied, and cells bound to the plate were stained with 0.006% crystal violet for 15 minutes at room temperature. Excess dye was rinsed away with distilled water. 300 μL of 100% DMSO was used to solubilize the dye that remained attached to cells. Microwell plates with Nunclon were used in this study (Thermo Fisher Scientific).

### Strain construction

*G. sulfurreducens* ∆*pgcA* was created using the markerless deletion method described in previously (Chan et al., 2015, Zacharoff et al., 2016). 1 kB up and downstream of GSU1761 (*pgcA*) was cloned into pk18mobsacB vector. This plasmid was mated into *G. sulfurreducens* via *E. coli* strain S17-1. The first round of selection was performed on (200 μg/mL) kanamycin NBFA plates to obtain recombinant cells. Kanamycin resistant colonies were restreaked and plated on 10% sucrose for a second round of selection for recombination events that resulted in reversion to wild type or gene deletion. Colonies from this round of selection were patched onto plates with and without kanamycin. Colonies sensitive to kanamycin were screened for loss of GSU1761 using PCR. GSU1761 was also cloned into pRK2-Geo2 (Chan et al., 2015) backbone for growth complementation testing. This plasmid contains a constitutive promoter from the *G. sulfurreducens* gene *acpP* (GSU1604). GSU1761 was also cloned into the pBAD202/D-TOPO® (Thermo Fisher Scientific) plasmid backbone which resulted in an arabinose inducible expression vector containing PgcA fused to a 6X – histidine tag on the carboxy terminus (pBAD202PgcA).

*S. oneidensis* was electroporated with pBAD202PgcA plasmid after passage of the expression plasmid through a methylation minus *E. coli* K12 ER2925 strain (New England Biolabs, Ipswich, MA). Transformants were selected on 50 μg/mL kanamycin infused LB plate. *S. oneidensis* was routinely cultured in lysogeny broth (LB) (Becton, Dickinson & co, Franklin Lakes, NJ). Plasmids and deletion strains were sequence confirmed via Sanger sequencing at UMGC, University of Minnesota. See Table 1 for strain designations.

**Table 1.** Strains used in this study

| Strain or Plasmid | Description | Source/Reference |
| --- | --- | --- |
| Strains |  |  |
| *G. sulfurreducens* PCA | Wild Type (ATCC 51573) | Lab collection |
| $\Delta pgcA$ *G. sulfurreducens* | Markerless deletion of *pgcA* gene in Wild Type *G. sulfurreducens* background | This study |
| *E. coli* S17-1 | Donor strain for conjugation | Simon, 1983 |
| *Shewanella oneidensis* | Wild Type | Myers and Nealson, 1988 |
| Plasmids |  |  |
| pK18mobsacB | Markerless deletion vector | Schafer, 1994 |
| pRK2-Geo2 | Vector control and backbone for complementation vector | Chan et al., 2015 |
| *ppgcA* | Complementation vector with constitutive expression of *pgcA*. *pgcA* was cloned into pRK2-Geo2. | This study |
| pBAD202/D-TOPO® | Arabinose inducible expression vector backbone | Thermo Fisher Scientific |
| pBAD202PgcA | Arabinose inducible expression vector containing PgcA | This study |

### Protein purification

10 mL cultures of the PgcA expressing strain of *S. oneidensis* were used to inoculate 1 liter of LB medium containing 50 μg/mL kanamycin. Cells were incubated at room temperature (25 °C) at slow rotation speed to achieve microaerobic conditions. The use of non-baffled shake flasks also decreased the amount of oxygen in the medium. The optical densities at 600 nm were monitored until an optical density of 0.5 was achieved. At this time 3 mM (final concentration) of arabinose was added to induce PgcA expression. 100 μM FeCl_3_ was also added at this time to increase the amount of bioavailable iron in the medium. Cells were pelleted 18 hours after induction at 4,000 × *g*. The pellet was washed with 100 mM Tris-HCl, 200 mM NaCl, pH 7.5 buffer. Resuspension and lysis via sonication (50% duty cycle, amplitude of 20%, 2 cm horn, for 30 minutes) was performed in the same buffer with lysozyme and DNase. Lysate was centrifuged at 30,000 × *g* for 30 minutes. The soluble fraction was loaded on to a nickel affinity column. Protein was eluted with 300 mM imidazole. Concentrated eluent was further purified with gel filtration or anion exchange chromatography. Gel filtration was done using a 45 cm length, 1 cm diameter column filled with Sepharose 6B (Sigma-Aldrich, St. Louis, MO). A flow rate of 1 mL/min was used during column equilibration and sample separation. Anion exchange separation was performed using HiTrap Q HP, 5 mL columns (GE Health Care, Uppsala, Sweden). A flow rate of 5 mL/min was used. Sample was loaded onto column in no salt 100 mM Tris-HCl. A gradient program was initiated using a mixture of 0.5 M NaCl, 100 mM Tris-HCl and no salt 100 mM Tris-HCl. Protein sample was monitored throughout purification using SDS-PAGE gel stained for total protein and for peroxidase activity based heme stain 3,3’,5,5’ tetramethylbenzidine (TMBZ) (Smith et al., 2015; Thomas et al., 1976).

### Mass spectrometry

Protein samples that resulted from nickel affinity purification were separated on a BisTris, SDS, 12.5% polyacrylamide gel. Bands at 41 kDa and 57 kDa were excised from the gel. Trypsin digest and LCMS mass spectrometry using Thermo LTQ were performed on each of the band sizes (Center for Mass Spectrometry and Proteomics, University of Minnesota). PEAKS Studio software was used to analyze fragments (BSI Informatics Solutions).

### Circular dichroism

Protein sample was dialyzed with 50 mM phosphate, ph 7.5, with 100 mM sodium fluoride to decrease background signal in the ultraviolet region (Greenfield, 2007). A JASCO-J815 spectropolarimeter was used to acquire circular dichroism spectra in the range of 185 nm to 600 nm. Samples were maintained at room temperature for the entirety of experimentation. Spectra were analyzed using K2D3 program (Pellegrini, 2015).

### Stimulation of Fe(III) oxide reduction by added PgcA

96 deep well plates were prepared with 20 mM Fe(III) oxide medium. Flavin mononucleotide (FMN) (0-200 μM), bovine serum albumin (BSA), horse heart cytochrome *c*, or purified PgcA (12 μM equivalent of each protein) were added prior to cell addition. 100 μL of 0.6 OD (600 nm) cells, either wild type *G. sulfurreducens*, or ∆*pgcA* strain, were mixed into the 1 ml wells. A negative control lacking cells was also included. Cells were allowed to reduce Fe(III) for 20 h in an anaerobic chamber with an atmosphere of 20% CO_2_, 75% N_2_, 5% H_2_. A FerroZine assay was used for Fe(II) quantification, as described above. Preliminary Fe(II) measurements conducted over 4h intervals verified that reduction was linear over this short incubation period.

### Sequences used for alignment of PgcA homologs

Sequences used for Figure 6 were obtained from (Strain, locus, GI number); *Geobacter sulfurreducens*, GSU1761, GI:637126441: *Geobacter uranirreducens*, Gura_0706, GI:640548206: *Geobacter bemidjiensis*, Gbem_1881, GI:642767873, *Geobacter sp. FRC-32*, Geob_3176, GI:643640481: *Geobacter sp. M21*, GM21_2329, GI:644869943: *Geobacter bremensis,* K419DRAFT_01717, GI:2524445678: *Geobacter argillaceus,* Ga0052872_00704, GI:2597449491: *Geobacter pickeringii,* Ga0069501_111509, GI:2633859152: *Desulfuromonas soudanensis WTL*, Ga0081808_112930, GI:2637110285: *Geobacter sulfurreducens AM-1*, Ga0098194_11, GI:2640720749: *Geobacter soli,* Ga0077628_111213, GI:2649969705: *Geobacter anodireducens,* Ga0133348_111806, GI:2689034555. After preliminary alignment by Clustal, sequences were trimmed to include only conserved repetitive/heme regions and re-aligned to obtain multifasta files as input for WebLogo3 using default parameters (Crooks et al, 2004).

## Results

### Predicted features of PgcA and related proteins

The amino acid sequence of *G. sulfurreducens* PgcA predicts three *c*–type heme binding (CXXCH) motifs separated by repetitive elements (Figure 1). The amino acids threonine (T) and proline (P) alternate to form a string of 29 PT_X_ repetitions, followed by a heme motif, and a second PT_x_-heme region (Figure 1). PT_x_-rich tandem repeats are found in many *G. sulfurreducens* relatives, while PA_x_-dominated repeats are found in strains such as *G. uraniireducens* and *Desulfuromonas soudanensis*. This general pattern could also be identified using PTRStalker to detect fuzzy tandem repeats (Pellegrini, 2015; Pellegrini et al., 2012), which detected many PgcA-like sequences in *Geobacter* genomes, and also predicted tandem repeats in extracellular cytochromes that did not contain PT_x_ or PA_x_ domains.

**Figure 1.**
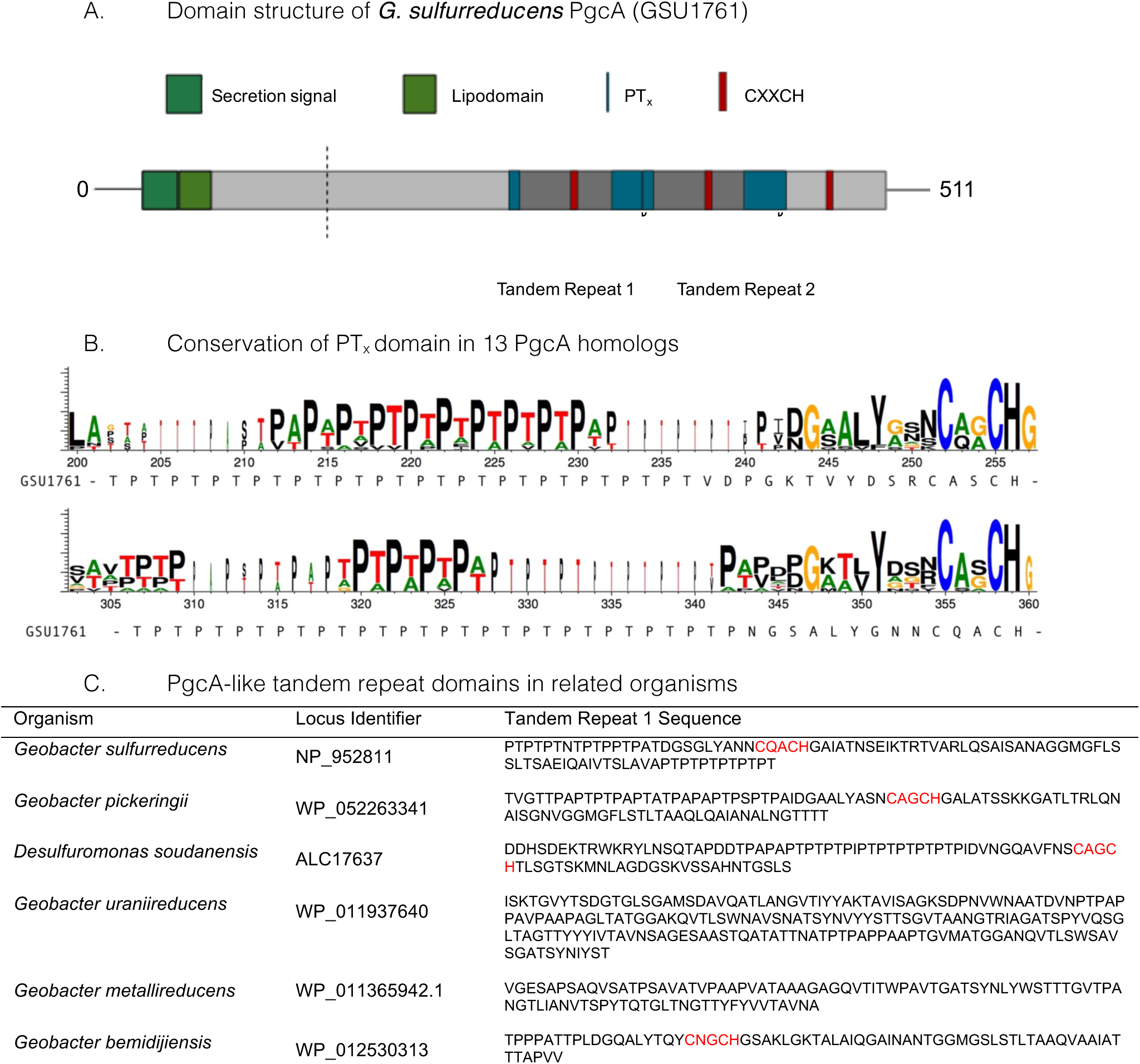
Characteristics of PgcA amino acid sequences. A. The full length protein includes a *sec-*secretion domain, lipid attachment domain (green), cleavage site identified by LC/MS, repetitive TPT_x_ domains (blue) and three CXXCH *c-* type cytochrome binding motifs (red). The tandem repeat of the PT_x_-CXXCH domain is highlighted. B. Conservation of repetitive TPT_x_ or APA_x_ pattern between heme domains in alignments of 13 homologs to PgcA. C. Other examples of repetitive domains identified by PTRStalker within putative secreted cytochromes of related strains.

The number of hemes within identified PgcA homologs varies. Only one CXXCH motif is observable in *G. metallireducens,* while six occur in *G. bemidjiensis*. The presence of PT_X_ repeats in the *G. sulfurreducens* sequence was notable, as Lower (Lower et al., 2008) found that proline in the tripeptide S/T-P-S/T restricts flexibility and positions serine/threonine hydroxyl groups for hydrogen bonding with metal oxide surfaces. Hematite association has also been observed near a short threonine-proline-serine motif near exposed heme groups in the *S. oneidensis* OmcA crystal structure (Edwards et al., 2012).

### Geobacter sulfurreducens cells lacking pgcA are Fe(III) oxide respiration deficient but capable of Fe(III) citrate and electrode respiration

A markerless mutant lacking *pcgA* showed no defect in reduction of the soluble electron acceptor Fe(III) citrate, and expression of *pgcA* via a constitutive promoter in ∆*pgcA* cells also had no effect on growth (Fig 2A). In contrast, when insoluble metals such as Fe(III) oxide (Figure 2B) or Mn(IV) oxide were present, reduction was severely impaired in the ∆*pgcA* strain. After 10 days of incubation, wild type *G. sulfurreducens* carrying an empty vector produced 36.3 mM Fe(II), while ∆*pgcA* produced only 9.0 mM Fe(II). Expression of *pgcA* from a constitutive promoter restored 75% of Fe(III) reduction activity, producing 27.4 mM Fe(II) in 10 days. A similar defect in Mn(IV) reduction was also observed in ∆*pgcA* mutants (data not shown).

**Figure 2.**
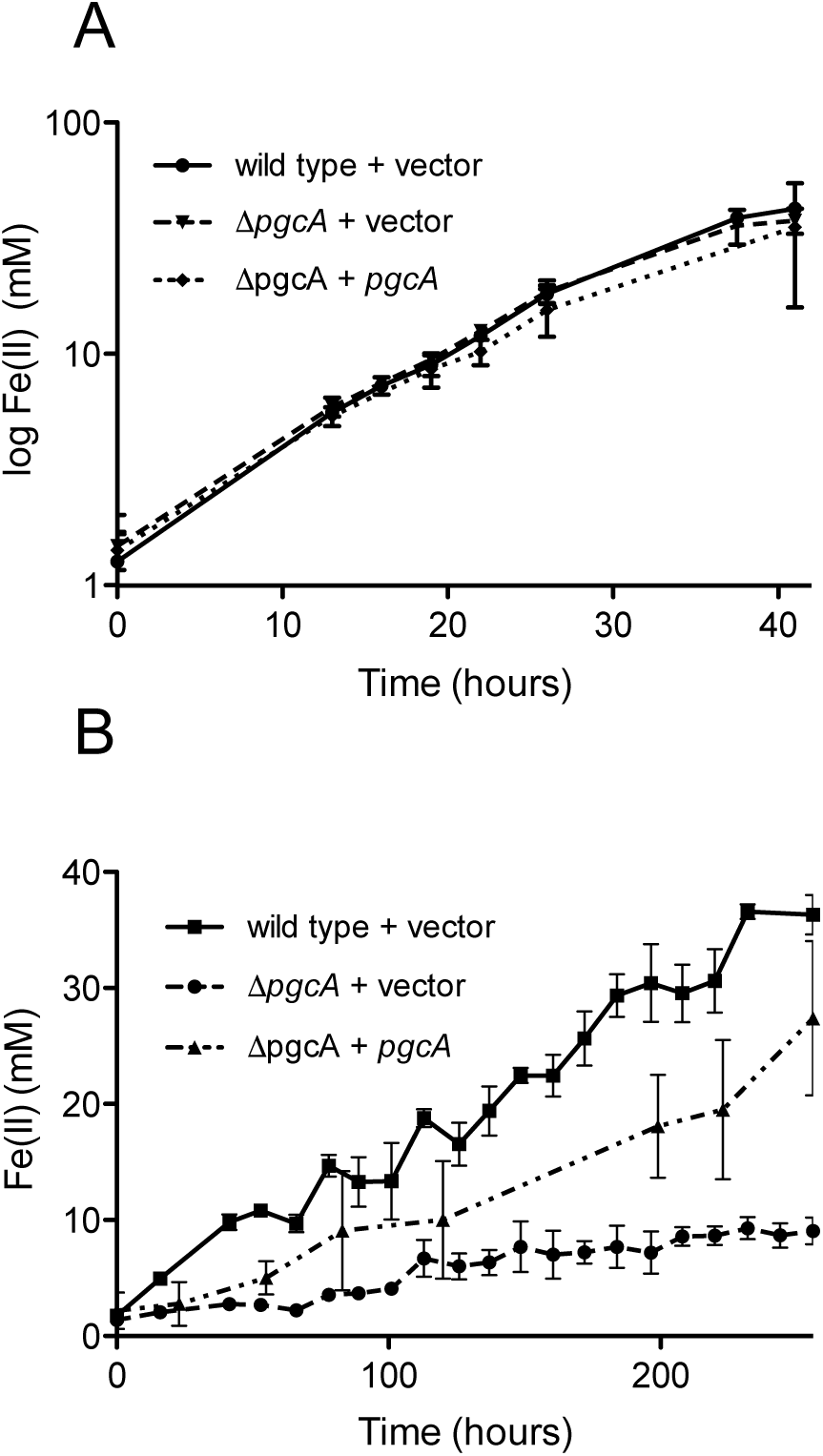
*G. sulfurreducens* mutants lacking *pgcA* are defective in reduction of insoluble Fe(III), but reduce soluble Fe(III) similar to wild type. A. Fe(III) citrate reduction by wild type carrying empty vector, ∆*pgcA* carrying empty vector, and ∆*pgcA* carrying the vector expressing *pgcA* from a constitutive promoter. B. Fe(III) oxide reduction by wild type carrying empty vector, ∆*pgcA* carrying empty vector, and ∆*pgcA* carrying the vector expressing *pgcA*. Error bars are +/- standard deviation of four replicates.

When cultivated using +0.24 V vs. SHE poised graphite electrodes as the electron acceptor, wild type and ∆*pgcA* cells demonstrated similar doubling times of 5.6 hours (n=3) vs. 5.5 hours (n=3) (Figure 3A). In addition, wild type and ∆*pgcA* cells reached a similar current density of 550 μA/cm^2^ within 3 days of growth. Complementation of ∆*pgcA in trans* also resulted in similar growth. Further evidence that PgcA played no role at any stage of electron transfer to electrodes was obtained from cyclic voltammetry scans over a wide (-0.4 V to +0.3 V) potential range, which were similar at all redox potentials. Similar results were obtained at −0.1 V vs. SHE, consistent with cyclic voltammetry data (Figure 3B).

**Figure 3.**
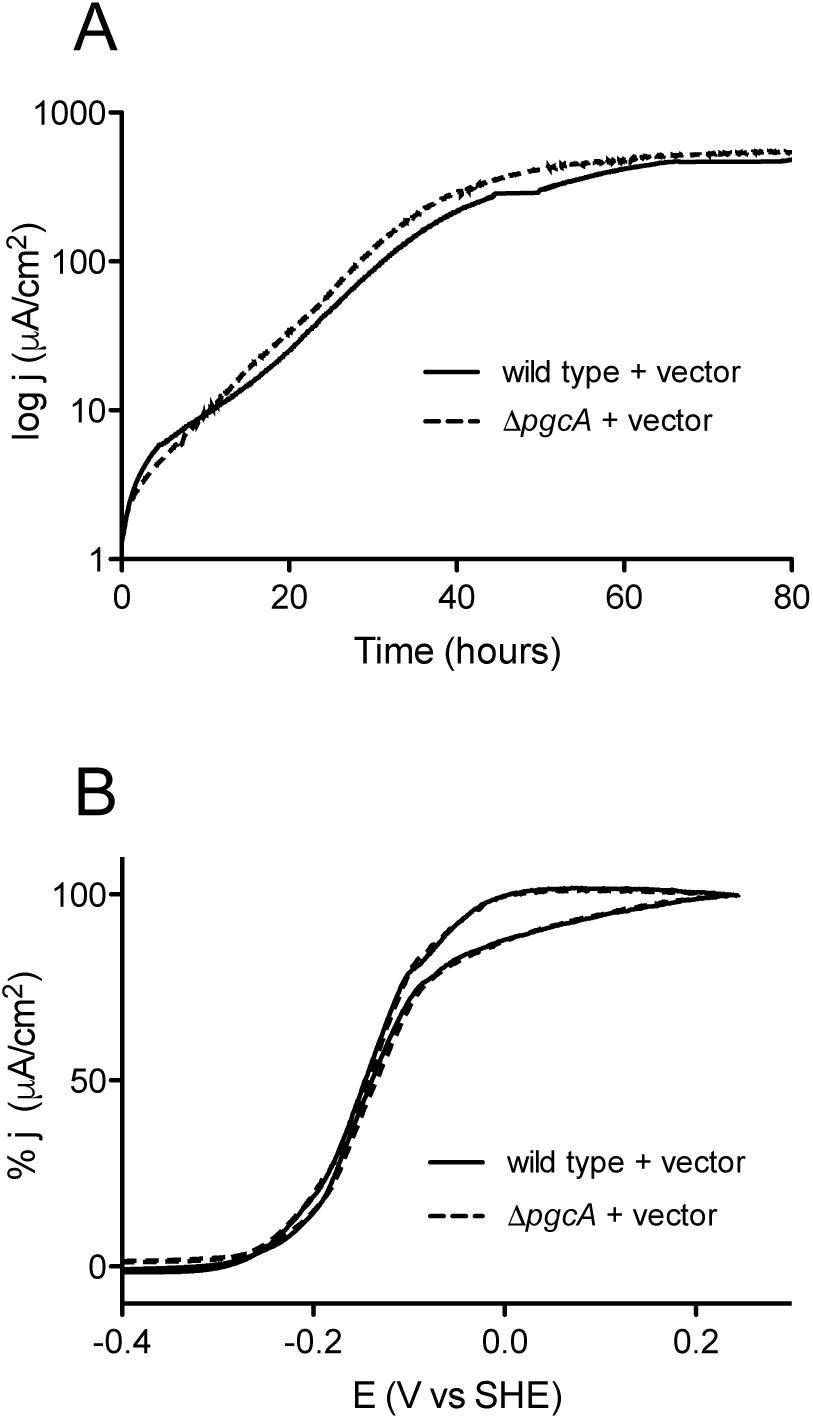
Deletion of *pgcA* does not affect growth of *G. sulfurreducens* on graphite electrodes. A. Working electrodes were poised at +0.24 mV vs. SHE, and current density, j, is expressed as μA/cm^2^ for wild type and ∆*pgcA* cells. B. Cyclic voltammetry of wild type cells compared to ∆*pgcA* mutants after 80 h of growth.

*Geobacter* strains lacking extracellular components can show increased binding to negatively charged surfaces, which has been correlated with defects in reduction of substrates. Mutants in the *xap* extracellular polysaccharide synthesis gene cluster show over 250% increases in attachment to negatively charged surfaces, and are also defective in binding poised graphite electrodes (Rollefson et al., 2009, 2011). In contrast, mutants showing 50-75% levels of attachment were not correlated with any reduction phenotypes. Binding was investigated in ∆*pgcA* cells grown to stationary phase with fumarate as the electron acceptor, and determined by a crystal violet attachment assay. Using polystyrene culture plates, ∆*pgcA* bound 79% as well as wild type, suggesting changes to the outer surface, but consistent with wild type-like interactions with electrodes.

### Biochemical assessment of PgcA

PgcA was expressed in *Shewanella oneidensis* under control of an arabinose-inducible promoter under microaerobic conditions, and successful incorporation of all three predicted hemes were determined by the pyridine hemochrome assay (Trumpower, 1987) and mass spectrometry. During purification, we consistently obtained both a large and small form; mass spectrometry of excised gel bands verified that these corresponded to 57 kDa and 41 kDa forms of PgcA, where the short form was truncated specifically at amino acid 127. This smaller variant was similar to the PgcA observed in evolved *G. sulfurreducens* KN400 strains overexpressing PgcA (Tremblay et al., 2011), as well as the unidentified cytochrome previously recovered from *G. sulfurreducens* PCA supernatants (Lloyd et al., 1999).

The visible spectrum of PgcA had an absorbance maximum at 405 nm in the oxidized state. The pyridine hemochrome assay extinction coefficient at 408 nm was 137,000 M^-1^cm^-1^, consistent with the incorporation of three hemes. PgcA was rapidly oxidized and reduced by ferricyanide and sodium dithionite, respectively. The reduced protein shifted to a maxima at 417 nm (γ or Soret), and demonstrated peaks at 518 nm (β), 552 nm (α). No additional changes in the absorbance spectrum, such as those characteristic of His-Met coordination, were observed in these experiments (Figure 4A).

**Figure 4.**
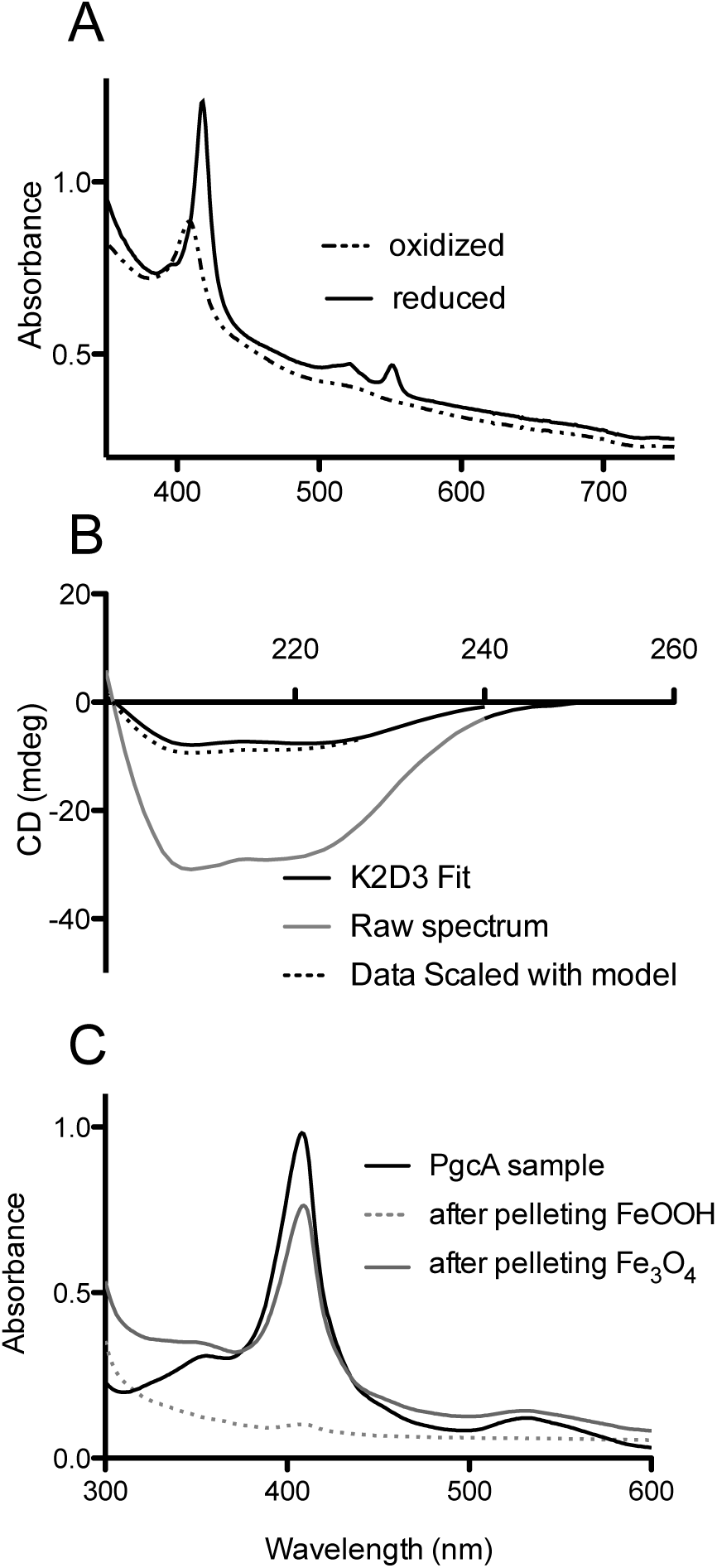
Biochemical characterization of *G. sulfurreducens* PgcA expressed in *Shewanella oneidensis*. Oxidized and reduced electronic absorption spectroscopy in the visible region. B. Circular dichroism of PgcA, using the 41 kDa processed form, modeled spectrum generated from K2D3 program. C. Spectroscopy of solutions after incubation and centrifugation of PgcA with oxidized Fe(III) oxide (dotted line) vs. magnetite (grey line).

Because of the significant amount of proline-rich repeats in PgcA, analyses of protein secondary structure was conducted using circular dichroism spectroscopy. Proline induced backbone rigidity can create unique secondary structures which alter regular alpha-helix/beta-sheet patterns, such as in the case of the collagen triplex helix. CD spectra were recorded in millidegrees (mdeg) from 200-240 nanometer wavelengths of fully oxidized, truncated (41 kDa) PgcA in pH 7.5 phosphate buffer with 100 mM sodium fluoride (Greenfield, 2007). No evidence of intrinsic disorder or unique secondary structures were detected in the experimental conditions (using K2D3). PgcA was composed of 70.5% alpha helical character and 5.2% beta-sheet character (Figure 4B), (Greenfield, 2007; Pellegrini, 2015). The alpha helix relative to beta sheet composition of PgcA was significantly more helical than the 10% alpha helix value estimated for OmcS (Qian et al., 2011) and the 13% value reported for OmcZ (Inoue et al., 2010), but is consistent with the alpha helical bias observed for other hemeproteins (Smith et al., 2010).

As some extracellular cytochromes (such as OmcS) show an affinity for binding Fe(III) oxides, PgcA was incubated with freshly prepared Fe(III) oxide, as well as biologically reduced magnetite at pH 6.5, where both of these minerals have a net positive charge (Kosmulski, 2011). Using absorbance at 410 nm to monitor soluble protein concentrations, PgcA showed an ability to bind the oxidized, but not reduced mineral. After incubation with Fe(III), all PgcA added to solution was removed by the pelleting of metal oxide particles. In contrast, incubation with magnetite removed less than 10% of PgcA from solution, and magnetite also did not reduce PgcA, based on spectroscopy. When BSA or horse heart cytochrome *c* were incubated with Fe(III) oxide or magnetite, both proteins remained in the supernatant (>90%).

### Purified PgcA added extracellularly can accelerate Fe(III) reduction capabilities of ∆pgcA cells to wild type levels

Purified PgcA was used to determine if PgcA added extracellularly could rescue the inability of ∆*pgcA* inability to reduce Fe(III), or accelerate activity in the wild type. As a control in these experiments, purified PgcA was compared with a known electron shuttle, flavin mononucleotide, as well as proteins not expected to facilitate electron transfer (bovine serum albumin and horse heart *c-*type cytochrome) (Hartshorne et al., 2007; Kotloski and Gralnick, 2013; Shi et al., 2012). All cells were pre-grown to a state of electron acceptor limitation, washed and incubated with Fe(III) oxide and acetate, and the accumulation of Fe(II) monitored for 20 hours.

Under these conditions, wild type *G. sulfurreducens* produced 3.4 mM Fe(II), while the ∆*pgcA* mutant only produced 0.13 mM Fe(II) (Figure 5). When purified PgcA was added, rates of Fe(III) reduction doubled in the wild type, but increased nearly 20-fold in the *pgcA* mutant. Addition of horse heart *c*-type cytochrome or bovine serum albumin at similar concentration had no stimulatory effect on either culture.

**Figure 5.**
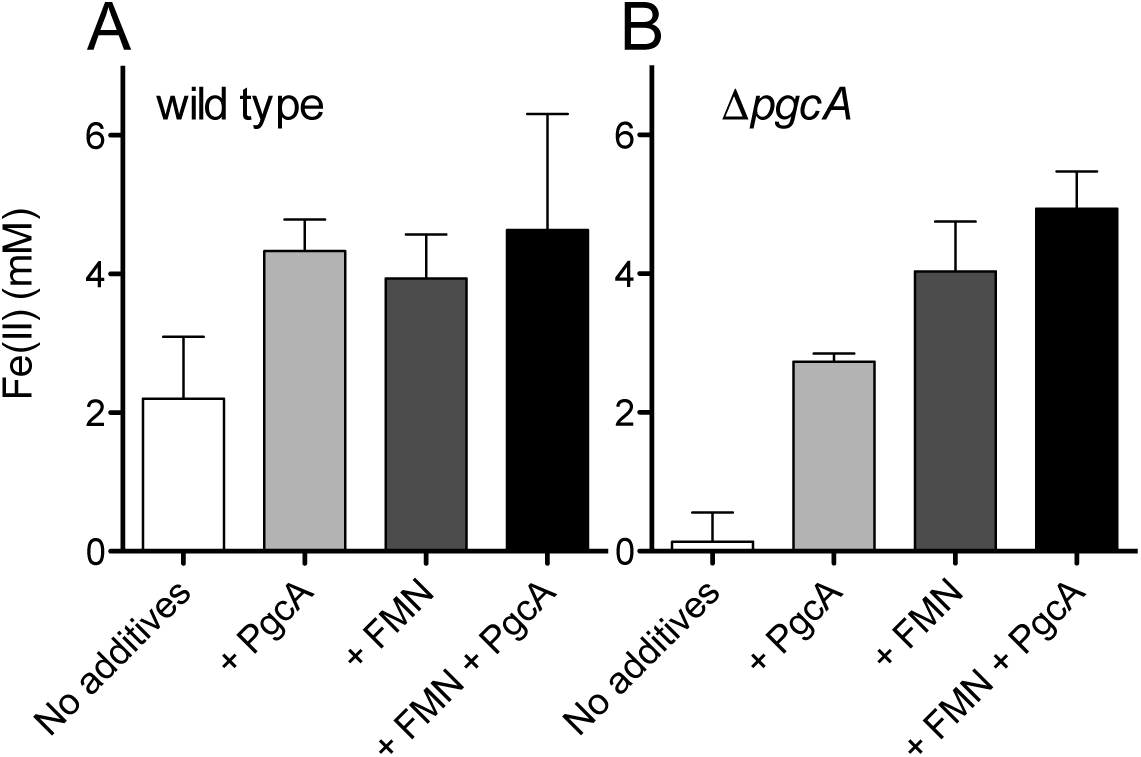
PgcA added to cell suspensions accelerates Fe(III) reduction. A. Washed wild type cells were incubated with Fe(III) oxide and acetate for 20 h. Cells were provided with 12μM PgcA, 50μM flavin mononucleotide (FMN), or both PgcA and FMN. B. Washed ∆*pgcA* cells incubated with Fe(III) oxide, 12μM PgcA, 50μM flavin mononucleotide (FMN), or both PgcA and FMN. Standard deviations are +/- 3 independent replicates.

When wild type cells were incubated with Fe(III) oxide and increasing amounts of flavin mononucleotide, rates of Fe(III) reduction improved until FMN concentrations reached 50 μM. Levels above 50 μM produced similar levels of stimulation to the wild type. Addition of 50 μM FMN accelerated metal reduction by the wild type cells similar to addition of PgcA, and stimulated reduction in ∆*pgcA* more than added PgcA alone. When both FMN and PgcA were added to the *pgcA* mutant, the effects were additive, resulting in the highest observed levels of stimulation, over 40-fold faster than the mutant alone.

## Discussion

The data presented here implicates a role for PgcA in electron transfer beyond the outer membrane, specifically during reduction of metal oxides compared to other acceptors such as electrodes or Fe(III) citrate. This role is consistent with studies correlating *pgcA* expression with Fe(III) oxide reduction, while the processing of PgcA explains repeated observations of a 41 kDa cytochrome in *Geobacter* supernatants. The protein has properties that support association with both cell surfaces and oxidized minerals, and in purified form, can be added extracellularly to rescue Fe(III) reduction by ∆*pgcA* mutants.

While PgcA can be recovered from cell supernatants, and soluble PgcA added to cell suspensions will accelerate metal reduction, the question remains whether it exists to diffuse freely between cells and metals, or is retained by flexible extracellular materials to increase the probability of cell-metal contacts. The strongest evidence arguing against soluble shuttle-like compounds in *G. sulfurreducens* is derived from experiments where metals entrapped in alginate beads are not reduced by cells. However, such beads are estimated to exclude proteins larger then 12 kDa, restricting PgcA from the encased iron (Nevin and Lovley, 2000).

Another way to examine whether PgcA could act as a freely soluble shuttle is to estimate the possible cost. To secrete enough PgcA to achieve a concentration of 10 μM in the space extending 1 μm in all directions from a cell 1 μm in diameter (an extracellular volume of 13.6 μm^3^) would require about 6.8 × 10^-15^ g protein. This would represent almost 7% of the 1 × 10^-13^ g protein in a *Geobacter* cell, a considerable cost (Figure 6). The high price of protein synthesis, combined with additional dilution of protein into the nearby environment, argues for mechanisms that keep proteins tethered to cells or functional at effective concentrations less than 1 μM, where the burden is calculated to be below 1% of cell protein.

**Figure 6.**
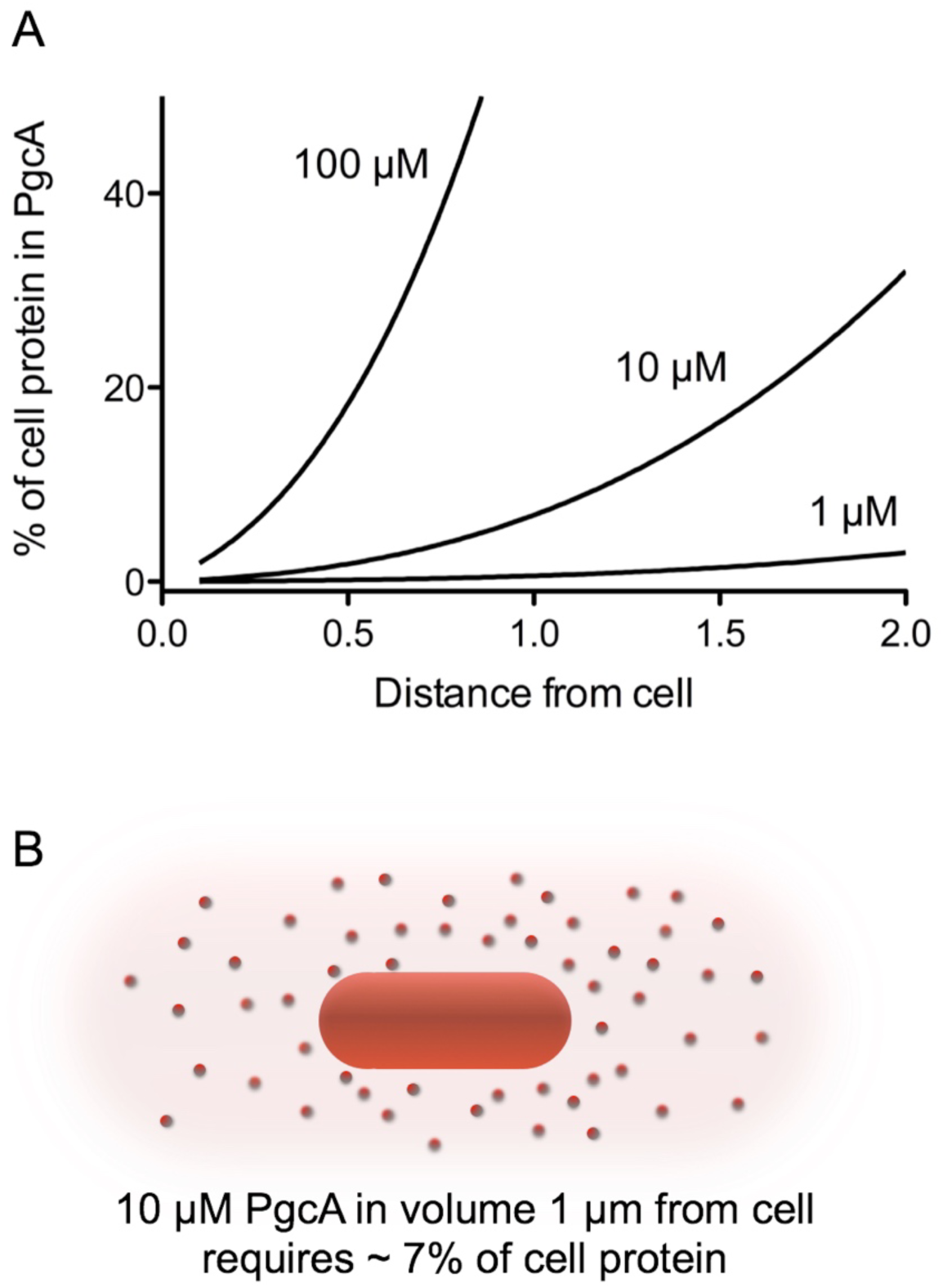
Calculated protein burden of using PgcA as a soluble shuttle across a range of concentrations and distances. A. The amount of a 50 kDa protein required to reach a given concentration in a given volume was calculated in grams, and expressed as a fraction of a standard 1 × 10^-13^ g *Geobacter* cell. B. Visualization of the volume around a cell needing to be filled; For simplicity calculations are based on a sphere extending from the cell membrane. A volume 1 μm in all directions is 13.6 μm^3^, while the volume 2 μm from a cell is 65 μm^3^. This exponential increase with distance rapidly increases the burden of a protein-based shuttling strategy.

The repetitive domains within PgcA also raise questions about localization. Tandem repeat domains are commonly associated with adhesion and biomineralization in secreted proteins (Paladin and Tosatto, 2015). Ice nucleation and antifreeze proteins contain simple TXT_x_ amino acid sequences (Kobashigawa et al., 2005), while TPT_X_ repeats of equal or greater length are found in secreted chitin binding, carbohydrate binding, and cellulose binding proteins. TPT_X_ repeats also occur in viral proteins of unknown function, including; *Thermoproteus tenax* virus isolated from a “sulfotaric mud hole” (Katti et al., 2000) and the ATV virus from *Acidianus convivator* (Prangishvili et al., 2006). The only function attributed to such repeats is based on the phage display work of Lower (Lower et al., 2008), who found S/T-P-S/T sequences bound metal oxides such as hematite. Such repeats exist as putative metal binding sites in OmcA/MtrC-family cytochromes, and are proposed to aid silica binding by silaffins in the alga *Thalassiosira pseudonana*. Based on our finding that PgcA bound Fe(III) oxides but not magnetite, one possibility is that the TPT_X_-rich region helps deliver the protein to oxidized metals, yet releases proteins as acceptors become reduced.

As components of the *Geobacter* electron transfer chain are revealed, a general theme of redox protein specialization has emerged. Different inner membrane cytochromes are required for reduction of low potential vs. high potential acceptors (Levar et al., 2014; Zacharoff et al., 2015), while in *S. oneidensis*, one inner membrane cytochrome is used for a range of metals, organic compounds, and electrodes (Gralnick and Newman, 2007; Marritt et al., 2012; Ross et al., 2011). Five outer membrane multiheme cytochrome conduits are functional in *G. sulfurreducens,* and only specific conduits and required for electrodes vs. Fe(III) and Mn(IV) oxides (Chan et al., 2017). In contrast, *Shewanella* uses only a single outer membrane complex for all acceptors (Gralnick and Newman, 2007). A similar diversity of secreted proteins with specific extracellular roles appears to be utilized by *G. sulfurreducens,* while no such secreted cytochromes have been found in *S. oneidensis.* One hypothesis is that the high reactivity and mobility of flavins produced by *S. oneidensis* act as a ‘universal translator’ between outer membrane cytochromes and minerals (Shi et al., 2012). In the absence of a redox-active shuttle, *Geobacter* may be forced to encode a wide assortment of secreted proteins such as PgcA to ensure direct electron transfer under all conditions.

## Acknowledgments

The Biophysical Resource Center at the University of Minnesota provided vital time and training on the JASCO-J815 circular dichroism spectropolarimeter. This study was supported by grant N000141612194 from the Office of Naval Research.

